# A model explaining environmental stiffness-dependent migration of fibroblasts with a focus on maturation of intracellular structures

**DOI:** 10.1101/2022.12.30.522353

**Authors:** Natsuki Saito, Tsubasa S Matsui, Daiki Matsunaga, Kana Furukawa, Shinji Deguchi

## Abstract

Cell migration is fundamental to many biological processes, while it remains elusive how cells modulate their migration upon different environmental stiffness. In this work, we focus on the structural maturity of actin stress fibers to explain the substrate stiffness-dependent emergence of different cell migration velocity. We demonstrate that fibroblasts migrate longer distances on softer elastic substrates, and the distance is increased by lowering the myosin-driven contractile force. Stress fibers, the major intracellular structure to generate and sustain contractile forces, were found to be less mature in structure on soft substrate than on stiff substrate. Based on these experimental results, we present a minimal mathematical model to capture the salient features of how the substrate stiffness alters the migration velocity. Specifically, the ability of cells to generate large contractile forces is limited on soft substrate according to the Hooke’s law. The inverse relationship between the cellular force and migration velocity is described by the Hill’s muscle equation. These mathematical descriptions suggest that the migration velocity is raised on softer substrate where cells exert a lower magnitude of contractile forces. Cells undergoing faster movement make stress fibers less mature in structure as mathematically described by the maturation model, thereby limiting the ability to sustain the force and in turn allowing for consistent increase in cell migration velocity on soft substrate again according to the Hooke’s law and Hill’s muscle equation, respectively. Thus, our model, reproducing the basic trend of the experimental results, provides insights into the mechanisms of environmental cue-dependent migratory behavior of cells.

## 1. Introduction

Cell migration is fundamental to many biological processes such as morphogenesis, wound healing, immune response, and tumor metastasis. During the migration process, cells sense environmental cues to behave adaptively in their surrounding milieu. Among the cues, substrate stiffness is increasingly attracting attention, in which particularly “durotaxis” has been a hot topic in recent years. Durotaxis is a directed cell migration where cells tend to move toward a stiffer region^1^. Many mathematical models have been proposed to describe this cellular response^2,3^, but the underlying mechanisms remain incompletely understood. Regarding the cellular response to substrate stiffness, cells are also known to alter their migration velocity at different substrate stiffness^4,5^. As cell migration is given birth to by the actin cytoskeleton, actin-based structures may take different states depending on the substrate stiffness to allow the cells to exhibit proper migration.

In this work, we focus on the role of the actin bundles or stress fibers in the adaptive behavior that cells modulate their migration upon different substrate stiffness. First, we demonstrate that fibroblasts migrate longer distances on softer substrates, and the distance is increased by lowering the myosin activity. As the actomyosin-driven contractile force is supposed to be smaller in magnitude on softer substrates given the principle of action– reaction, these results suggest that the cellular contractile force inhibits migration. We further show that the actomyosin-based stress fibers, as a force generator, are less stable on soft substrates, suggesting that the structural maturity is less advanced. Based on these results, we build a mathematical model to describe the salient features of how the substrate stiffness alters the migration velocity. Our model captures the basic trend of the experimental results and thus provides insights into the mechanisms of complex cell migration.

## 2. Materials and Method

### 2.1 Cell culture and transection

Human foreskin fibroblasts HFF-1 (ATCC) were cultured with DMEM (high glucose) including L-glutamine and phenol red (Wako) supplemented with 15% fetal bovine serum (Sigma-Aldrich) and 1% penicillin-streptomycin solution (Wako) in a humidified 5% CO^2^ incubator at 37°C. The plasmids encoding human myosin regulatory light chain (MRLC)^6^ gene combined with EGFP were transfected to cells using Lipofectamine LTX Reagent with PLUS Reagent (Thermo Fisher Scientific) according to the manufacturer’s instruction.

### 2.2 Elastic substrates

A prepolymer solution of polydimethylsiloxane (PDMS) was prepared at 10:1, 20:1, and 80:1 w/w ratio of the base polymer to the cross-linker (Sylgard 184, Dow Corning) to endow the PDMS substrate with different values of stiffness. The silicone elastomer component was coated on a plastic dish or glass-bottom dish using a spin coater (Kyowa Riken) at 1500 rpm, baked at 60°C for 20 h to cure the gel, and then treated with oxygen plasma (5 mA, 60 sec, SEDE-GE, Meiwafosis). The Young’s modulus of the PDMS substrate was evaluated by putting fluorescent microbeads onto the substrate to measure the resulting indentation with confocal microscopy and analysis based on the Hertz contact theory^7^ (Supporting information). The Young’s modulus of the 10:1, 20:1, and 80:1 substrate was determined to be 1.2 MPa, 490 kPa, and 1.4 kPa, respectively.

### 2.3 Cell imaging

Cells stained with CellTracker™ Green CMFDA (Invitrogen) were cultured for live cell imaging. Images were acquired with phase-contrast microscopy (IX-83, Olympus) equipped with a humidified 5% CO^2^ stage incubator (Tokai Hit) for 24 h at a 5-min interval. Time-lapse images were analyzed by using ImageJ/Fiji software (NIH). The position of individual cells was detected with the fluorescent marker to calculate the total migration distance by binarizing the images with a threshold value and obtaining the center of the fluorescent cells. To investigate the role of cellular contractile force in cell migration, (-)-blebbistatin (5 or 10 µM, Wako) was applied to cells to inhibit the activity of myosin II. An equivalent amount of dimethyl sulfoxide was applied to cells as vehicle control.

### 2.5 Fluorescence recovery after photobleaching

Cells expressing EGFP-MRLC were cultured on a glass-bottom dish (φ27 mm, No. 1S, #3970-035, Iwaki) or PDMS (10:1) dish in a humidified 5% CO^2^ stage incubator at 37°C. Fluorescence recovery after photobleaching (FRAP) experiments were performed by confocal microscopy (FV1000, Olympus) with a 60x oil immersion objective lens (NA = 1.42). Photobleaching was induced using 405/440-nm-wavelength lasers on EGFP-MRLC-labeled stress fibers. Images were taken at a 5-sec interval for 4 min.

### 2.6 Statistical analysis

Statistical analysis was performed using Origin 2021 software. Differences were calculated based on the Mann-Whitney U-test for variables with a non-Gaussian distribution or the unpaired Student’s T-test for variables with a Gaussian distribution, with a significance level of p < 0.05 (*) or p < 0.01 (**).

### 2.7 Model analysis

We propose a minimal model that describes a substrate stiffness-dependent migration of fibroblasts. Based on our observations that cell migration is enhanced on soft substrate compared to stiff substrate as described below, we assume that the state of stress fibers, contributing to substrate stiffness sensing, changes with the substrate stiffness. Mechanical force generated in a material is described in general by the product of its extensional rigidity (i.e., the product of the cross-sectional area and Young’s modulus in 1-dimensional cases) and effective deformation, or strain, measured from its non-stressed state according to the Hooke’s law. The extensional rigidity of individual stress fibers is comparable to or higher than that of the substrate sustaining the tension generated in stress fibers^8^. In such springs connected in series, the weaker one is dominant. Therefore, the tension in stress fibers as well as in the serially connected structure, *F*(*t*), is described by

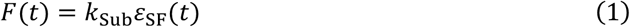

where *k*_Sub_, *ε*_SF_(*t*), and *t* are the stiffness of the substrate or more exactly extensional rigidity, effective strain of stress fibers, and time, respectively. The effective strain represents the ability of stress fibers to be physically able to sustain the tension *F*(*t*), which is determined based on the structural maturity of the stress fibers as described below.

Recent experiments indicate that cellular response to mechanical loading is qualitatively similar to that of muscle cells in a manner called Hill’s equation^9^. The mean contractile velocity of stress fibers *µ* is accordingly described by

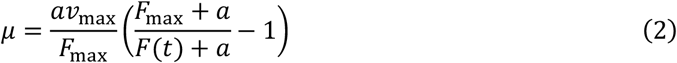

where *a, F*_max_, and *ν*_max_ are a coefficient, the maximum tension of stress fibers, and the maximum contractile velocity at *F* = 0, respectively. Here, *a* and *F*_max_ are 6 ×10^−8^ N and 0.1 N, respectively, both of which are determined to allow stress fibers to sustain a tension of 50 nN at a velocity of ∼0.3 μm/min and 100 nN at ∼0.2 μm/min according to previous experiments^10,11^. *ν*_max_ is 0.6 μm/min determined from the maximum velocity observed in the present experiments. The contractile velocity of stress fibers *ν*_SF_(*t*) is stochastically determined from a normal distribution with the mean *µ* and standard deviation *σ* described as

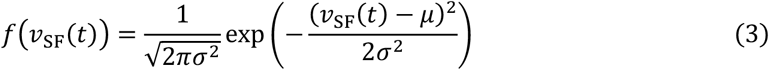

where *σ* is assumed to be a constant of 0.1 (μm/min), which is determined to have the same order as that observed in the present experiments. Unlike motile cells where dynamic stress fibers are utilized for migration, more stable stress fibers in nonmotile cells including HFF-1 fibroblasts are utilized for substrate stiffness sensing rather than for migration^12,13^. In terms of cell signaling, RhoA and Rac1, which activate stress fibers and lamellipodia, respectively, are competing with each other^14^. Thus, the activation of stable stress fibers and resulting stabilization and immobilization are supposed to suppress the lamellipodia-mediated cell migration. To capture this feature in our model, the magnitude of cell migration *ν*_cell_(*t*) is determined by the state of stress fibers as

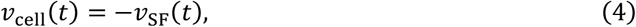

in which the negative sign is given because the direction is the opposite between the contraction of stress fibers and migration of cells. Here, Eq. (4) contains only a coefficient of 1 other than the negative sign because the relative quantitative relationship is already considered in *a, F*_max_, and *ν*_max_ in Eq. (2).

While generating tension, stress fibers are subject to turnover, by which their constituent molecules are constantly exchanged. In relatively dynamic stress fibers that are contracting faster than more static ones bearing larger forces, turnover is supposed to occur more frequently to eventually be less mature in structure^12,13^. The extent of the structural maturity is evaluated by an immobile fraction determined in fluorescence recovery after photobleaching (FRAP) experiments, and a first-order single exponential model or maturation model is often used for FRAP analysis. Given these features, the extent of maturation or maturation index *M*(*t*) is defined as a function of the magnitude of the contractile velocity of stress fibers, namely

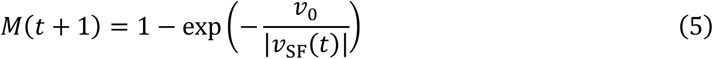

where *ν*_0_ is a constant of 0.3 μm/min, which is determined to capture the tendency of the immobile fraction observed in FRAP experiments. Here, time *t* is incremented by 1, and the stress fiber velocity determines the extent of maturation of the next time step so that the time is shifted by 1 between them. The extent of maturation determines the effective strain, or the ability of stress fibers to be able to generate and sustain the tension, so that

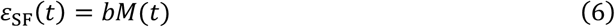

where *b* is a constant of 0.1, which is determined according to previous experiments where stress fibers possess a maximum strain of approximately 0.1^10,11^. The total migration distance *L*_total._ is determined by the product of the magnitude of instantaneous cell velocity |*ν*_cell_(*t*)| and time interval Δ*t*, namely

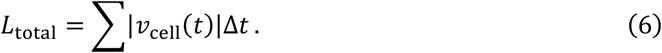

Using this model, *L*_total._ is numerically computed with a change in substrate stiffness *k*_Sub_. To mimic the effect of the myosin inhibitor (-)-blebbistatin in this model, the tension *F*(*t*) is reduced by multiplying by a factor of 0.1–1.0.

## 3. Results

### 3.1 Cells migrate longer distances on softer substrate

To explore the effect of changing the substrate stiffness on cell migration velocity, HFF-1 fibroblasts were plated on the substrate with the different levels of stiffness, i.e., 10:1 (Young’s modulus of 1.20 MPa), 20:1 (490 kPa), and 80:1 (1.45 kPa). The trajectory of cells during migration is analyzed to determine the total distance reached over the course of 1 day (Fig.1a). The distance was increased in a linear manner over time and was greater on softer substrate (Fig.1b).

**Figure 1.**
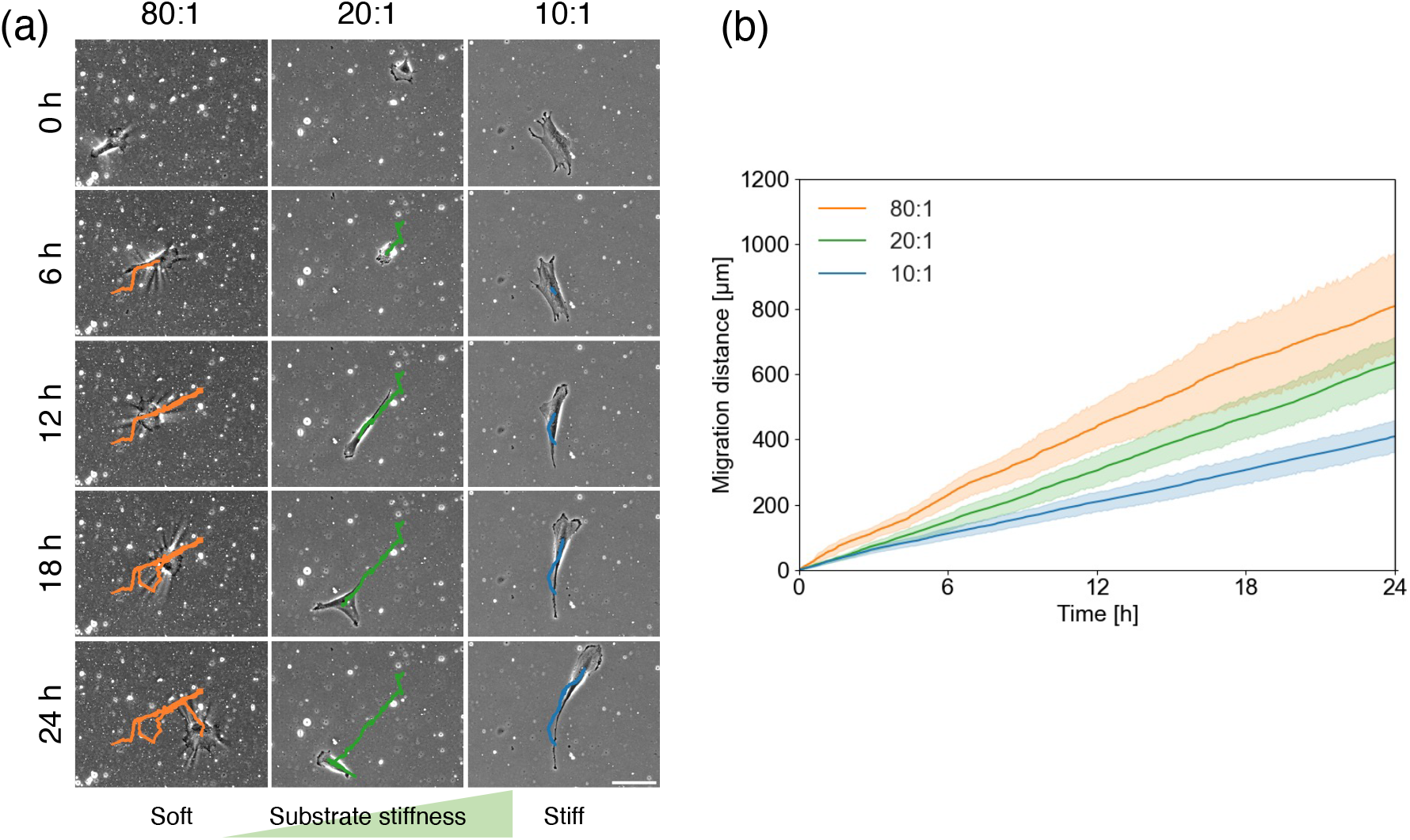
Experimental results of cell migration. (a) Time-series of the trajectories of HFF-1 cells migrating for 24 h on the substrate with a different stiffness. (b) Total migration distance of cells plated on substrates with different levels of stiffness. 80:1 (*n* = 10 cells), 20:1 (*n* = 18 cells), and 10:1 (*n* = 20 cells). Light shaded bar represents 95% confidence interval. Data are from at least 3 independent experiments. Scale, 200 µm

### 3.2 Cellular contractile force suppresses migration

To explore the effect of decreasing contractile force on cell migration, the total migration distance was analyzed with different concentrations of (-)-blebbistatin, an inhibitor of the inorganic phosphate release in the actomyosin ATPase and in turn of the cellular contractile force generation. To evaluate the relative change caused by the force inhibition, the migration distance fis normalized as

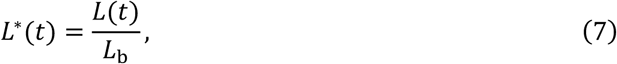

where *L*(*t*) and *L*_b_ are the migration distance at an arbitrary time *t* and that just before cells are treated with (-)-blebbistatin, respectively. Specifically, cells were treated with (-)-blebbistatin or its vehicle at 6 h after the observation began. While no significant difference was observed at 6 h after the treatment with 5-μM (-)-blebbistatin compared to control, the 10-μM treatment caused a significant increase in the migration distance (Fig. 2a, b). Thus, the cellular force and migration velocity were found to be inversely related to each other, consistent with the behavior described by Eqs. (2)–(4) with the Hill’s muscle equation. This inverse relationship seems to be independent of the choice of drugs because it has been reported that decrease in cellular force induced by many types of drugs has a tendency to cause increase in migratory potential^15^. While linear temporal increase in migration distance was observed in control as well as 5 μM, the 10-μM treatment ends up nonlinear increase over time probably because it requires some time to take effect. Indeed, the increase became almost linear after sufficient time of about 1 h.

**Figure 2.**
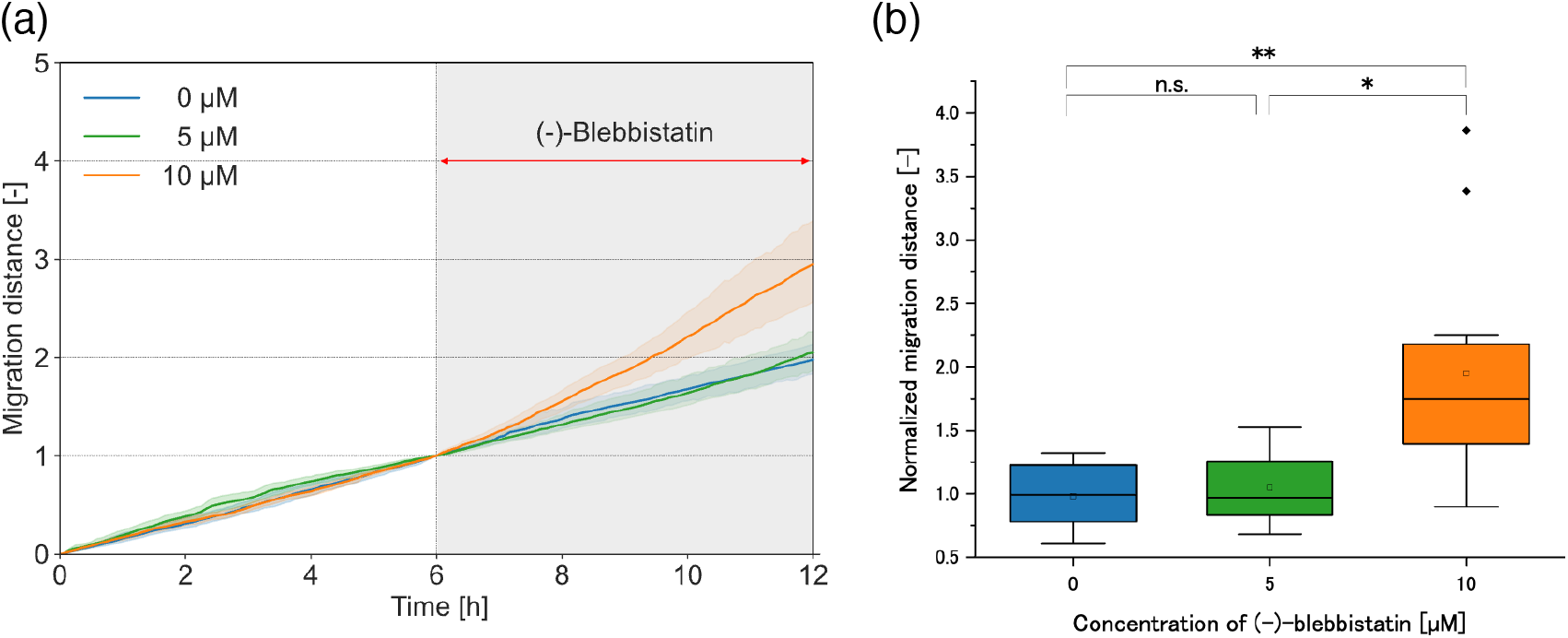
The effect of (-)-blebbistatin on the migration distance of cells. (a) Total migration distance of cells plated on 10:1 substrate. The distance was normalized by a value at 6 h. Cells were treated 6h after the observation with (-)-blebbistatin at a concentration of 0 µM (DMSO, *n* = 10 cells), 5 µM (*n* = 7 cells), and 10 µM (*n* = 14 cells). Light shaded bar represents 95% confidence interval. (b) Migration distance that cells reached within 6 h after (-)-blebbistatin treatment was evaluated at 12 h after the initiation of observation. Boxes represent the 25th and 75th percentiles and the median. Open squares represent the means. Whiskers extend from the box ends to the most remote points excluding the outliers defined as 1.5× the interquartile range. Data are from at least 2 independent experiments.

### 3.3 Stress fibers tend to be more mature in structure on stiffer substrate

We analyzed how the difference in substrate stiffness affects the turnover, or the extent of the stability, of stress fibers by conducting FRAP experiments. To represent stress fibers, one of the major constituents MRLC^16,17^ was targeted for measurement (Fig.3a). We found in preliminary experiments that, with soft substrates such as 20:1, it was hard to focus on the whole individual stress fibers because a nonnegligible level of out-of-plane deformations occurs on the top of the substrate, making it difficult to perform FRAP experiments. For this technical reason, only 10:1 was used here as a soft substrate, while a glass substrate was instead chosen as a stiff substrate representative. Note that these two substrates, 10:1 and glass, are not coated in advance with extracellular matrix but are made hydrophilic by oxygen plasma treatment, thus possessing similar chemical properties. While 10:1 is 1.20 MPa in Young’s modulus, the glass-bottom dish is known to be several tens of GPa and thus be stiffer compared to the former^18^. Upon photobleaching, MRLC molecules on the stiff (glass) substrate recovered more slowly than ones on the soft (10:1) substrate (Fig. 3b), indicating that there are larger amounts of unexchanged MRLC molecules on the stiff substrate. This tendency was quantified by analyzing the ratio of the fluorescence recovery at the end timepoint *t* = 235 s (Fig. 3c), suggesting that stress fibers in cells plated on stiffer substrate are more stable and mature.

**Figure 3.**
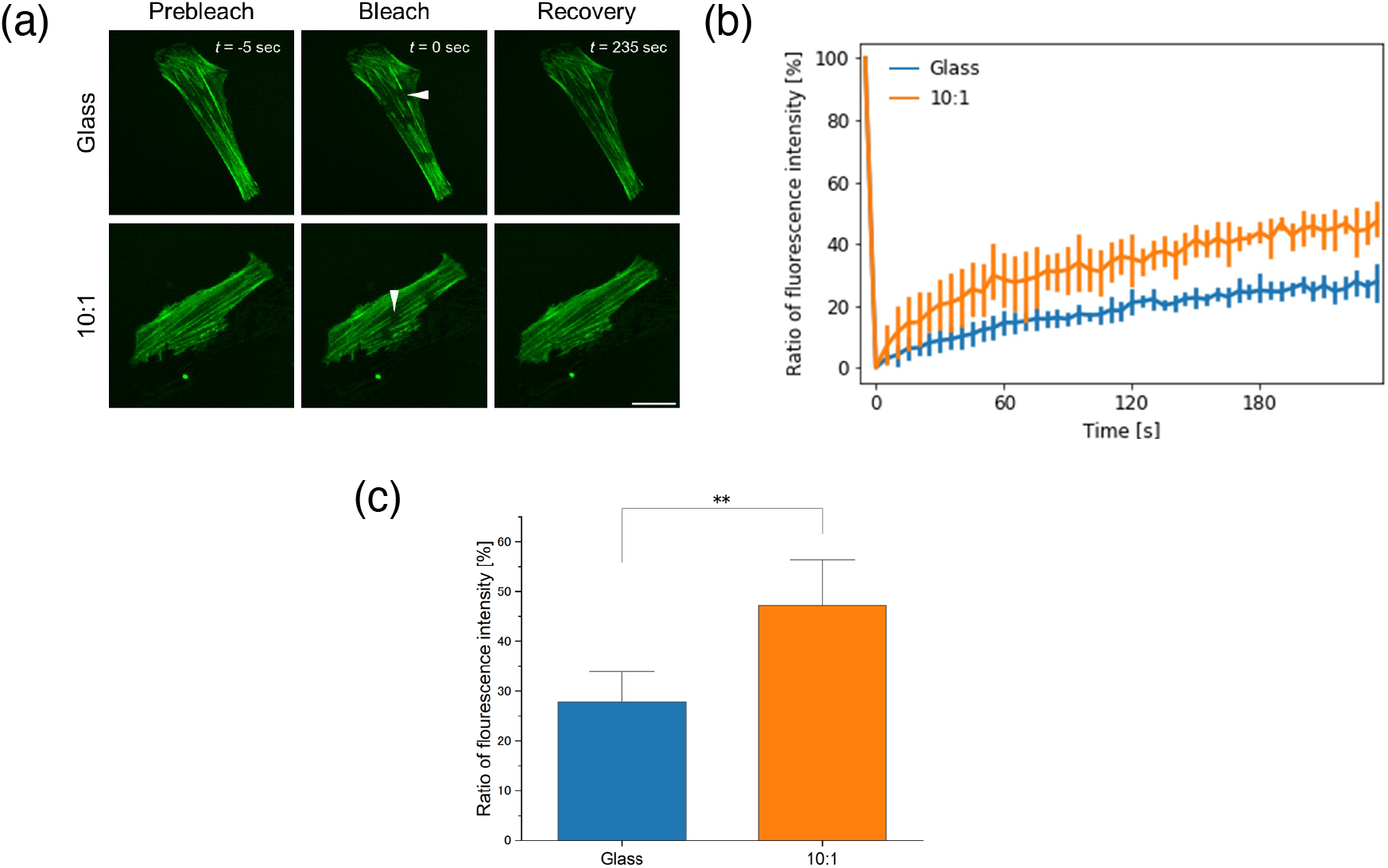
Evaluating the maturity of stress fibers. (a) FRAP experiments were performed on HFF-1 cells expressing EGFP-MRLC on glass or 10:1 substrate. Arrowhead indicates the bleached area at *t* = 0. While photobleaching was sometimes done at multiple regions, only one individual stress fiber was selected, from one individual cell, as a sample for FRAP analysis in the aim of capturing the typical behavior of the cell sample. Scale, 20 µm. (b) Time-series change in fluorescence intensity, in which all the values are normalized between the initial and photo-bleached states and are shown in percentage. Blue and orange show stiff (glass) and soft (10:1) substrate, respectively. Vertical bar represents 95% confidence interval. (c) Ratio of fluorescence intensity at *t* = 235 s showed that cells on the soft substrate exchange their molecules more rapidly than those on the stiff substrate. Data are from *N* = 3 independent experiments.

### 3.4 The model-based simulation captures the features of cell migration

We built a minimal model to describe the essence of the substrate stiffness-dependent cell migration. In the simulation, we give the difference only in stiffness of the substrate *k*_Sub_. The total length achieved by the cell migration is increased in a linear manner over time by lowering *k*_Sub_ (Fig. 4a), thus consistent with the experimental results (Fig. 1b). Next, we also simulated the (-)-blebbistatin treatment experiment (Fig. 2a) by allowing the cells to migrate with the same parameter sets and then reducing, at *t* = 6 h and for the rest of the time, the tension *F*(*t*) with a factor of 0.1, 0.4, 0.7, or 1.0. As expected, longer migration distance was reached with a lower level of tension (Fig. 4b), thus again consistent with the experimental results (Fig. 2a).

**Figure 4.**
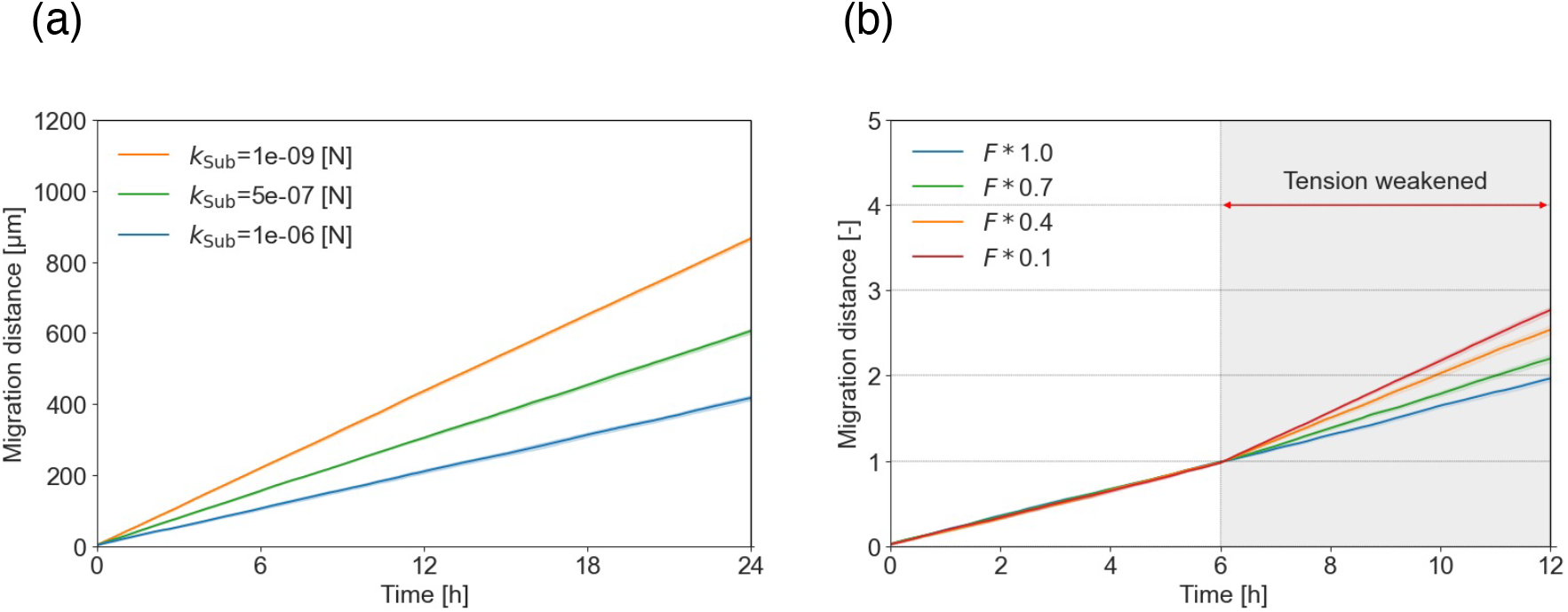
Simulation results of the effect of different substrate stiffness on cell migration distance. (a) Results with *k*_Sub_ =1 × 10^−9^, 5 × 10^−7^, and 1 × 10^−6^ are shown, which mimic the actual behaviors of 80:1, 20:1, and 10:1, respectively, shown in Fig. 1b. (b) The behavior of cells treated with (-)-blebbistatin, shown in Fig. 2a, is mimicked by reducing the magnitude of *F*(*t*) by a factor 0.1, 0.4, 0.7, and 1.0, ending up within the range of migration distance observed in the experiments.

**Figure 5.**
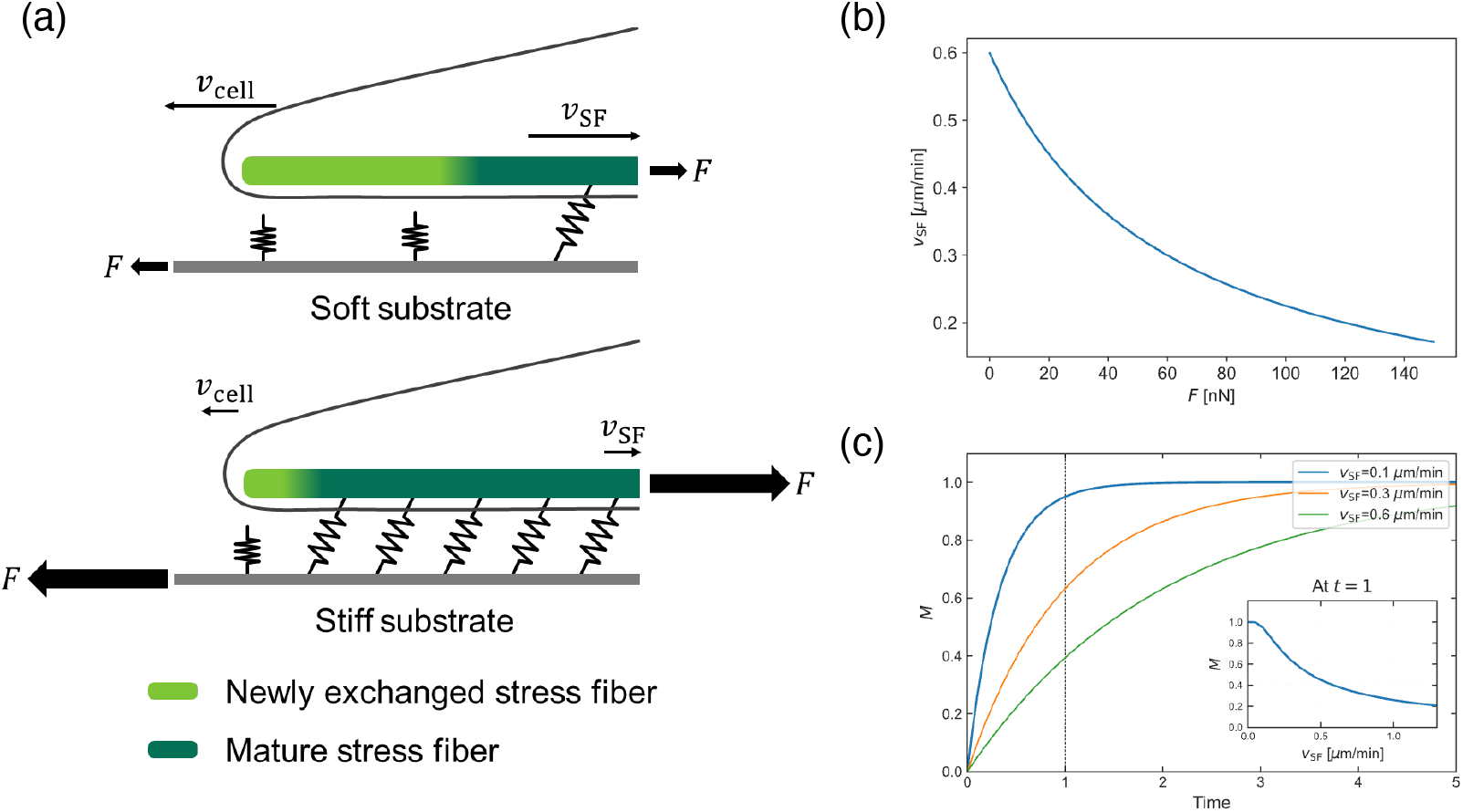
Conceptual illustration of the minimal model of fibroblast migration. (a) Cells on stiff substrate (below) exert a larger magnitude of contractile tension *F* (depicted by the large arrows) compared to the case of soft substrate (top) owing to the more mature stress fibers (dark green), resulting in slower stress fiber velocity *ν*_SF_ as well as cell migration velocity *ν*_SF._ as depicted by the short arrows. Here, the larger number of the springs in the case below represents a higher level of substrate stiffness. The tension generated in stress fibers is better sustained together with the underlying substrate in the case below because of the more advanced maturation of stress fibers as illustrated by the firmer connection between the intracellular and extracellular structures. (b) The relationship between *ν*_SF_ and *F*, plotted according to Eq. (2), i.e., Hill’s equation, which is one of the key factors of the model and underlies the reduced stress fiber velocity and cell motility upon increased contractile force. (c) The effect of changing the contractile velocity of stress fibers *ν*_SF_ on the time-series of Maturation index *M*, plotted according to Eq. (5), i.e., maturation model. The inset shows the relationship between *M* and *ν*_SF_ at *t* = 1 (dotted line). This motility-dependent extent of the structural maturation is another key factor of the model and underlies the advanced force transmission among the intracellular and extracellular structures on stiff substrate.

## 4. Discussion

In this work, we propose a mathematical model describing the salient features of the substrate stiffness-dependent migration of fibroblasts based on the experimental observations on cells and stress fibers. Stress fibers play a critical role in generating cellular contractile force, which in turn affects lamellipodia that facilitate cell migration via the competitive RhoA–Rac1 crosstalk^19–21^. Our experiments suggest that stiffer substrates slow cell migration compared to softer ones in a contractile force-dependent manner (Figs. 1 and 2). Thus, the cellular force and migratory potential are inversely related, suggesting that cells behave in a manner consistent with the behavior described by Eqs. (2)–(4) including the Hill’s muscle equation. Moreover, FRAP experiments revealed that stress fibers in cells plated on the stiff substrate are more prone to be structurally mature than those on the soft substrate, suggesting that stiff substrate stabilizes the intracellular force generator (Fig. 3). We extracted its features in the minimal model by focusing on the mechanical environment-dependent extent of the maturity of the intracellular structures associated with cell migration (Fig. 4). Specifically, stress fibers become more mature on stiff substrate rather than on soft substrate, strengthening the ability of exerting large contractile force onto the underlying substrate. The increased force is inherently accompanied by decrease in cell motility as well as in contractile velocity of stress fibers^22^. Collectively, the relatively mature stress fibers on stiff substrate slow cell migration.

Numerous mechanistic models have been proposed to describe cell migration. Focusing on ones that consider the interaction with the surrounding environment, synchronized beating of cardiomyocytes^23,24^ and circumvention of interference between motile keratocytes^23,24^ have been analyzed by modeling stress fibers as a force dipole. Focusing further on the effect of substrate stiffness on more normal proliferative cells such as fibroblasts used in the present study, cells were approximated to be composed of a set of springs and dampers mimicking the actin cytoskeleton^2,3^, and then cell movement was predicted from force balance. Cellular tendency of migrating within identical directions on stiff substrate has been analyzed using a random walk model^25,26^, but the biological origin of the stiffness-driven persistency of the migration direction is not clearly described. Viscoelastic interaction between cells and substrates has been analyzed to explain cell migration^27^, but the involvement of the actin cytoskeleton is not explicitly considered. In other studies, motor-clutch models^28,29^ that consider force transmission among the actomyosin structure, cell adhesions, and substrates have explained the cell behavior on elastic substrates. The role of focal adhesions as a mechanosensor^30,31^ and the involvement of both mechanics and chemistry^32^ in substrate stiffness-dependent cell migration have also been analyzed. Nevertheless, the qualitative change in stress fibers intrinsically induced by the different environmental stiffness was not clearly described in these studies, while adaptive remodeling achieved by continuous turnover is a salient feature of the intracellular structure^22^. In this regard, the central idea of our model is the inclusion of the structural maturity of stress fibers, which is not explicitly documented in the previous studies on environmental stiffness-dependent cell migration and is therefore discussed in more details below.

We accounted for the origin of the stiffness-dependent behavior by distinguishing the extent of the structural maturity of stress fibers. We considered that the maturation affects the effective strain to be physically able to sustain contractile forces. As mechanical force in a material is inherently determined by the product of strain and stiffness or more accurately extensional rigidity, we expressed it according to the Hooke’s law (Eq. 1). The force then determines the velocity of cell migration according to the Hill’s muscle equation as the macroscopic consequence is the same as that of muscles, in which larger forces lead to slower motion (Eq. 2). Here, the knowledge of the RhoA–Rac1 relationship underlies this modeling, which implies better development of stress fibers is followed by less activation of lamellipodia^20,21^. This relationship is indeed consistent with our observations that at a higher stiffness cells migrate more slowly, and the reduction in contractile force results in increase in migration (Figs. 1 and 2). The correlation in velocity between stress fibers and cells is expressed as a stochastic process following the Gaussian distribution (Eqs. 3 and 4). The stress fibers undergoing a high level of cell migration remain relatively immature in structure to thereby reduce their effective potential for sustaining forces on the substrate (Eq. 5, Fig. 3). The above scheme of the Hooke’s law then comes again to eventually enable the stiffness-dependent cell migration (Eqs. 1 and 6). Consequently, the output of our model was in good agreement with the experimental results (Fig. 4).

The contribution of the maturity of cell–substrate adhesions to the cellular response to the surrounding environment has been extensively studied^33,34^. In contrast, it remains poorly understood how the extent of stress fiber maturity is altered in a cell context-dependent manner to regulate cell migration. Ventral stress fibers in fibroblasts are known to be comprised of at least 135 different proteins, and 63 of them are altered in expression only by repeating cell passage to induce senescence^16^. Such specific molecular identification underlying the stress fiber maturation is beyond the scope of the present study, but we instead captured the salient features with the minimal physicochemical model by introducing the factor of maturity level based on the FRAP observations. Thus, while the present model is a conceptual one highlighting the importance of the extent of intracellular structural maturity in the complicated cell processes, it was found to be enough to simulate the experimental results. Note again that our present focus is placed on characterizing the minimal elements that allow cells to exhibit a different migration velocity depending on the environmental stiffness, and therefore additional elements will be necessary to associate our model with durotaxis. Nevertheless, our analysis provides fresh insights particularly regarding the involvement of structural maturation of the intracellular components for better understanding of the whole complicated mechanism.

## Acknowledgment

We thank Sho Yokoyama for providing a part of the stiffness data and Rie Taniguchi for technical support.

## Author contributions

Conceptualization, N.S., S.D.; Data Curation, S.D.; Formal Analysis, N.S.; Funding Acquisition, S.D.; Investigation, N.S.; Methodology, N.S., T.S.M., S.D.; Project Administration, S.D.; Resources, T.S.M., S.D.; Supervision, S.D.; Validation, N.S., T.S.M., D.M., K.F., S.D.; Visualization, N.S., D.M., K.F., S.D.; Writing, N.S., S.D.

## Supporting information

The Young’s modulus of the PDMS substrate of 10:1, 30:1, 50:1, and 80:1 was measured according to the method described previously (Yokoyama et al., 2017). We then obtained a regression line using the least-square method (Fig. S1), in which “[Substrate stiffness (kPa)] = 15877 x [PDMS mixture ratio] – 264.85.” Here, the mixture ratio is calculated as, for example in the case of 10:1, 1/(10 + 1) = 0.0909. The experiments using a PDMS mixture ratio of 0.0476 (= 1/(20 + 1)), namely 20:1, were finally adopted in this study, we determined its Young’s modulus from the regression line to be 490 kPa. These values are quantitatively similar to, or at least in the same order as, data shown in a previous report measured using a tensile tester on the same but bulk types of PDMS (Prager-Khoutorsky et al., 2011) as shown in Fig. S2, in which 20:1 is supposed to be 868 kPa in Young’s modulus. Our estimate on the Young’s modulus of 20:1 (namely 490 kPa) seems thus reasonable, while we particularly focus in the present study on the qualitative feature of the substrate stiffness-dependent cell migration, and the value is not necessarily critical to our results.

**Fig. S1.**
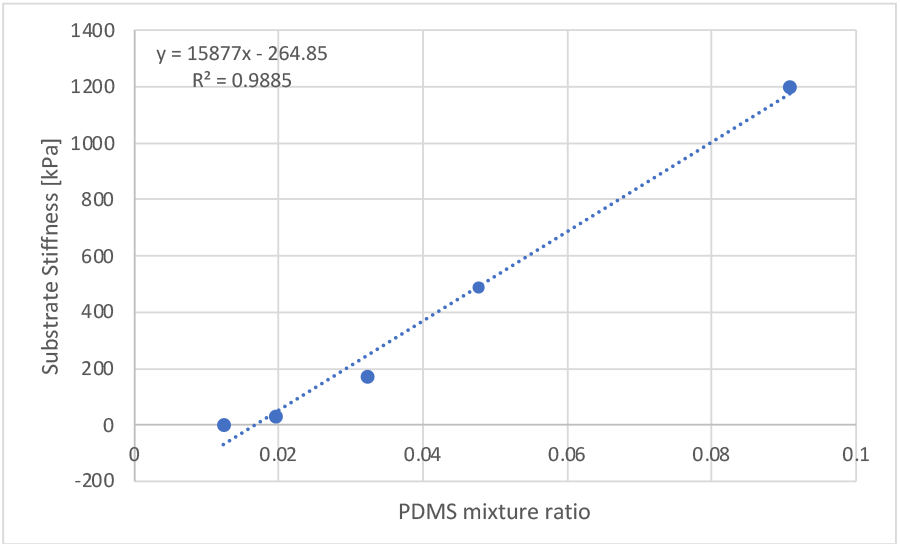
Our experimental data

**Fig. S2.**
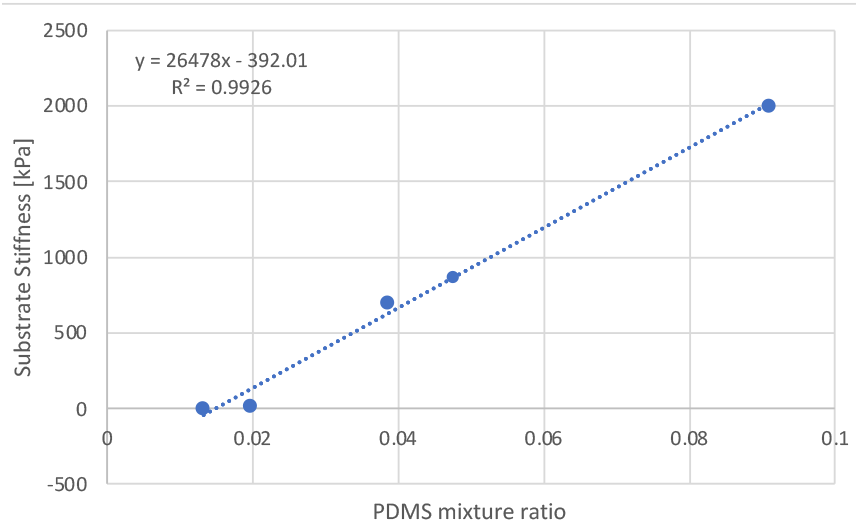
Data from Prager-Khoutorsky et al.

